# Extensive field-sampling reveals the uniqueness of a trophy mountain goat population

**DOI:** 10.1101/484592

**Authors:** Jessica Breen, Meghan Britt, Justin B. Johnson, Daria Martchenko, Yasaman Shakeri, Boyd Porter, Kevin S. White, Aaron B.A. Shafer

**Author notes:** Contributed equally (Jessica Breen, Meghan Britt, Justin B. Johnson). Trent University, Peterborough, Ontario Canada, K9J 7B8, *Email (Aaron B.A. Shafer).

## Abstract

Collaborations between academic researchers and agencies is crucial for published genetic research to have a tangible impact on conservation and wildlife management. Such partnerships are particularly important for elusive species where the difficult terrain requires that a significant amount of resources and a combination of methods be used to estimate population parameters needed for conservation and management. We report a multi-year academic-agency collaboration on the North American mountain goat that used an extensive field-sampling of genetic and phenotypic data to determine whether, and to what degree, genetic and phenotypic differences separate an isolated population of mountain goats on the Cleveland Peninsula from those in southeast Alaska. We observed significantly larger horns on the peninsula and the population appears demographically isolated. Isolation-by-distance accompanied by limited migration and low effective population size on the Cleveland Peninsula suggest this population will continue to lose genetic diversity. While the large horns of mountain goats have generated interest in re-opening mountain goat harvest on Cleveland Peninsula, our genetic data suggest this population is vulnerable to demographic and environmental perturbations and is unlikely to support a sustained harvest.

---

The relevance of genetic data to conservation and management has been questioned in the academic literature. Vernesi et al. (2008) observed that few researchers used their data or methods to suggest practical solutions, and Bowman et al. (2016) showed that most evolution and genetics studies were under-applied compared to ecology. An essays by Hoban et al. (2013) raised similar concerns, arguing for more collaboration between academics and practitioners, while meta-analyses revealed a positive relationship between the inclusion of non-academic co-authors and linking data to policy or management recommendations (Britt et al. 2018), notwithstanding a general long-term decline in implementation of research to management and conservation (Mair et al. 2018). Despite the gap between geneticists and on-the-ground practitioners (Shafer et al. 2015, Haig et al. 2016), headway for genetically-informed management recommendations in the peer-reviewed literature can be achieved via partnering with non-academics and non-biologists (Britt et al. 2018, Mair et al. 2018). The benefit presumably stems from the sharing of non-overlapping resources, expertise, and knowledge.

The purchase of hunting licences in the United States grosses close to one billion dollars annually (US Fish And Wildlife Service 2017). The economic benefits derived from hunting often mean significant resources are made available to management and conservation of wildlife that are guided by the principles of the North American Model of Wildlife Conservation (Organ et al. 2012). In Alaska, approximately 500 mountain goats (*Oreamnos americanus*) are harvested each year with the majority of hunting concentrated in coastal areas. The mountain goat is characterized by a thick white coat with relatively short, black horns that have been used by indigenous groups for centuries and are sought after by hunters (Festa-Bianchet and Côté 2007). As such, the sustained persistence of healthy mountain goat populations entails both societal gain and enduring cultural significance.

The mountain goat’s extreme habitat and sensitivity to anthropogenic disturbance poses a significant challenge relative to other species in terms of population monitoring (White et al. 2011) and sightability models are often implemented (Poole 2007, Rice et al. 2009). As a result, a combination of observational, GPS tracking, and genetic data have been used to fill in knowledge gaps regarding population patterns and processes (Mainguy et al. 2009, Shafer et al. 2012a). Molecular genetics, specifically non-invasive genetic monitoring methods, represents an important tool for obtaining accurate and cost-effective population data that is less invasive for the target species (Taberlet and Luikart 1999). Genetic tools allow for delineating populations boundaries and estimating parameters such as migration rates (i.e. gene flow), heterozygosity, and effective population size (*N*_e_), all of which can inform management decisions. This is relevant to Alaska mountain goats as they are managed at the hunt unit level, not at the game management unit or GMU (see Fig. 1). Hunt units are defined based on expert opinion and available data like survey data and harvest vital rates.

**Figure 1.**
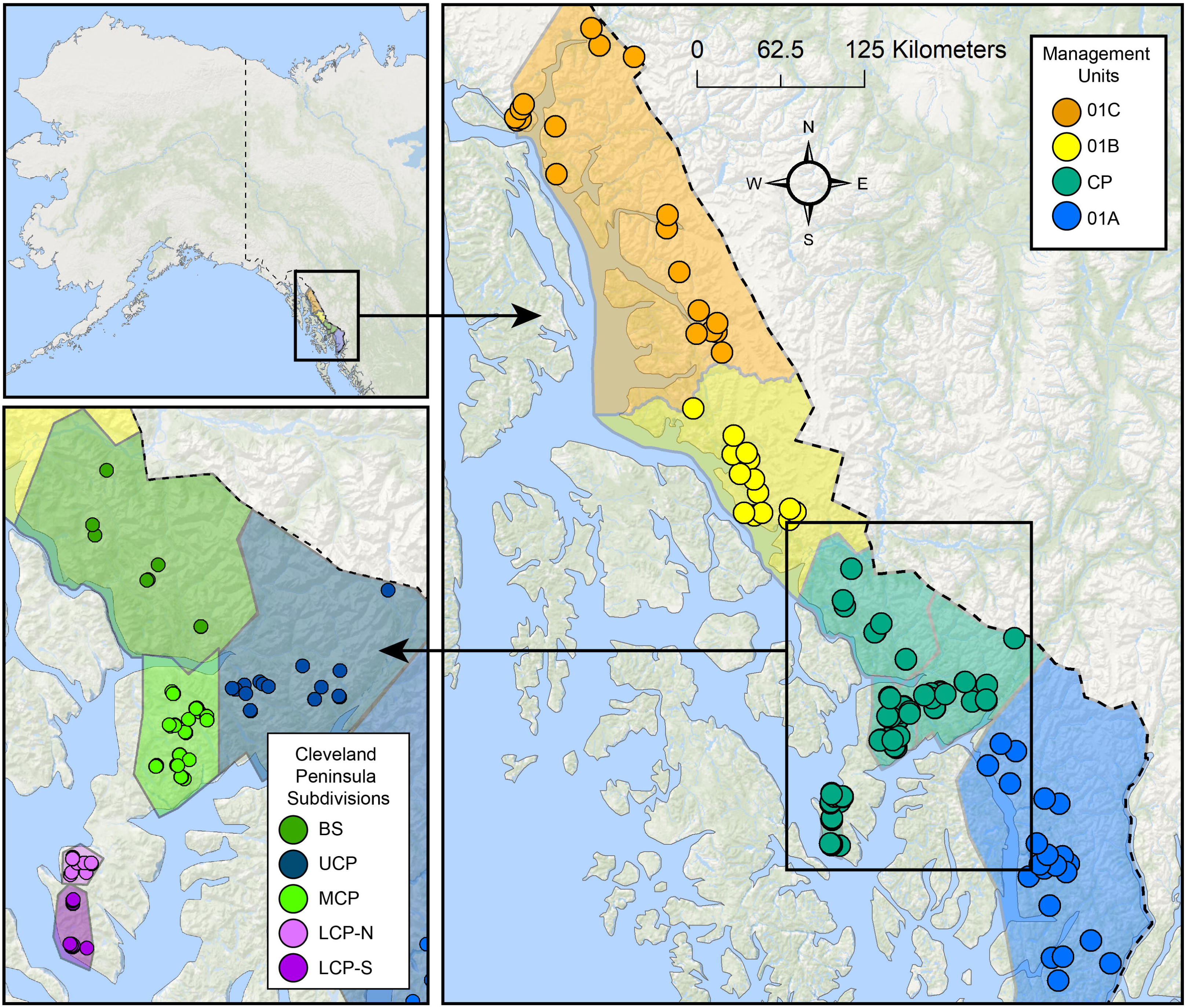
Map showing management unit GMU 01 in southeast Alaska and the Cleveland Peninsula (a); map of the six sub-divisions in the Cleveland Peninsula (CP); Lower Cleveland Peninsula North (LCP-N), Lower Cleveland Peninsula South (LCP-S), Middle Cleveland Peninsula (MCP), Upper Cleveland Peninsula (UCP) and Bradfield-Stikine (BS)

The Cleveland Peninsula (Fig. 1) is an interesting case study involving mountain goat management as it is separated from the main mountain range by 32 km of low elevation terrain that is generally deemed unsuitable for the species (White et al. 2010). There are additional management and societal interests in the population due to disproportionately large horns, with four of the top ten Boone and Crockett records coming from the peninsula (Buckner et al. 2009; Boyd Porter, unpublished data). Based on the strict habitat requirements of mountain goats and the apparent geographic isolation, it is currently unclear whether there is any mountain goat movement between the population(s) on the peninsula and those inhabiting the Coast Mountains in southeast Alaska.

In the 1980s there were an estimated 50-70 individuals across 9 suitable habitat patches found on the lower part of the Cleveland Peninsula (Smith and Raedeke 1982). In 2004, high harvests and an apparent population decline led managers to close the area to hunting. In 2013-2014, only 4 patches were occupied with an estimated 40 individuals (K. S. White, unpublished data). As little is known about the population(s) on the peninsula, more information is required to guide management decisions, especially as it pertains to harvest. Specifically, knowledge about the degree to which the population(s) are isolated and how much genetic interchange is occurring is needed to assess the resilience of the population to stochastic events and the extent to which genetic rescue could aid the population in the event of natural or anthropogenic mediated declines. Here, we present a multiyear academic-agency collaboration that used an extensive field sampling effort combining genetic and phenotypic data to determine whether, and to what degree, genetic and phenotypic differences separate the population(s) on the Cleveland Peninsula from those found in southeast Alaska.

## METHODS

### Sample Collection

A pilot project was initiated in 2009 by the Alaska Department of Fish and Game (ADFG) to obtain precise population estimates for the Lower Cleveland Peninsula, entailing the capture of 13 mountain goats. Mountain goats were fitted with radio-collars, phenotypic measurements were taken, and ear punches were obtained for DNA analysis. Tissue samples from mountain goats harvested from GMU 01 between 2005-2015 were also used for this analysis (Fig. 1).

In the summers of 2014, 2016, 2017, and 2018, helicopter crews identified mountain goats on the Cleveland Peninsula and recorded their locations. Where individuals were sighted, a search was conducted on foot for fecal pellets. Non-invasive pellet samples were directly swabbed and placed in Longmire’s solution (100 mM Tris, 100 mM EDTA, 10 mM NaCl, 0.5 % SDS, 0.2 % sodium azide). Samples from the Cleveland Peninsula were divided into four sampling regions based on the geographic location of major ridges; Lower Cleveland Peninsula North (LCP-N), Lower Cleveland Peninsula South (LCP-S), Middle Cleveland Peninsula (MCP), Upper Cleveland Peninsula (UCP). A transition zone Bradfield-Stikine (B-S) between the mainland was sampled, along with neighbouring GMUs 01A, 01B, and 01C (see Fig. 1).

### Horn measurements

Horn measurements from mountain goats captured by ADFG in southeastern Alaska between 2005-2017 were collated and compared to captured mountain goats from the LCP. We evaluated horn growth between males in southeast Alaska by first generating length-by-age regression curves. A polynomial regression was fit with horn length as the response variable and location and age as explanatory variables, with a quadratic function applied to the age variable. We compared the annuli length at year 1 and 2 using a Student’s t-test (note only the first two years were available for LCP). Analyses were conducted in R v. 3.4.1.

### DNA Extraction and Genotyping

Genomic DNA was extracted from the tissue and the non-invasive fecal and hair samples with QIAGEN DNeasy Blood and Tissue Kits (Qiagen, Inc., Valencia, CA). Fecal samples were swabbed with a cotton swab that had been wetted with de-ionized water and then placed downward in a 1.5mL tube. Ear punches and swabs that were collected in the field were placed wholly into 1.5mL tubes. All samples were digested overnight (~24 hours) in 180uL of Qiagen’s Buffer ATL and 20uL of Pro-K before they were extracted following standard Qiagen protocol. Extractions were quantified using a Qubit 3.0 Fluorometer (ThermoFisher Scientific) and stored at −20° C.

A total of 18 polymorphic microsatellite loci were amplified in 3 PCR pools (Table S1). All non-invasive samples were amplified in triplicate. Each polymerase chain reaction followed the approach of Shafer et al. (2010), and samples were genotyped on an ABI 3730 DNA Analyzer and scored using the program Geneious R10 (Kearse et al. 2012). Non-invasive sample genotype calls for each locus were compared across three replicates and if there were disagreements (i.e. alleles were not the same), the peaks were compared visually, and the weak ones removed (RFU < 250). If more than one replicate at a marker had discrepancy between heterozygosity or homozygosity after three replicates, the marker was considered missing (scored 0). The resulting Cleveland Peninsula dataset was screened with the R package *Allelematch 2.03* (Galpern et al. 2012) to remove potential duplicate individuals. Individual samples and microsatellite loci with over 50% missing data were removed from analysis.

The number of alleles, heterozygosity, inbreeding and differentiation indices were determined using the program GenAlEx v6.5 (Peakall and Smouse 2006). We estimated *N*_m_, or the effective number of migrants per generation between the sampled populations. We compared diversity statistics between the tissue and non-invasive samples from the LCP-S to determine whether sample quality influenced the results. All of our samples were collected within 15 years (1-3 generations) thereby reflecting a snapshot of population history; as such we estimated the effective population size (*N*_e_) using a single point in time and linkage disequilibrium method available in NeEstimator V2.1 (Do et al. 2014). Isolation-by-distance was conducted by taking the natural logarithm of the Euclidean distance between sampling area centroids and comparing this to *F*_ST_. Significance was assessed using a Mantel test and 1,000 permutations (Mantel 1967).

The Bayesian clustering program STRUCTURE v.2.3.4 was then used to identify subpopulations (Pritchard et al. 2000). Admixture models within STRUCTURE were parallelized with StrAuto for K = 1-10 (i.e. number of clusters) with a burn-in of 5×10^5^ followed by 1 × 10^6^ MCMCs with 20 replicate runs for each K. The results of the runs for each K were sorted and averaged with CLUMPP 1.1.2 (Jakobsson and Rosenberg 2007). This process was completed once with the PopFlag parameter set to 1, which defines *a priori* population assignments for known groups, and once with the PopFlag parameter set to 0. The assignment results were comparable with and without the PopFlag parameter, so we used our results without predefined reference groups to allow the software to find its own population divisions. We also used a principal component analysis (PCA) in the R package *adegenet* (Jombart 2008) to visualize population genetic structure and confirm the STRUCTURE results.

### Mitochondrial Sequencing and Analysis

All samples from the LCP were sequenced at the mitochondrial DNA d-loop to confirm species identity as the population is sympatric with Sitka black-tailed deer (*Odocoileus hemionus sitkensis*). A subset of MCP, UCP, and B-S samples were sequenced to include in the analysis. Samples were amplified following the approach and primers in Shafer et al. (2011) with negative controls. The PCR product was run on a gel to ensure sufficient DNA, and then cleaned via ExoSAP-IT protocol (ThermoFisher Scientific) and ethanol precipitation prior to submission. Samples were sequenced on an ABI 3730 DNA Analyzer, and edited and aligned with the program BioEdit (Hall 1999). For mtDNA analysis, 38 additional *Oreamnos americanus* d-loop sequences from southeast Alaska were obtained from GenBank to create a spatial outgroup. We quantified the number of unique haplotypes on the Cleveland Peninsula and plotted this using a haplotype network.

## Results

A total of 419 horn measurements were acquired and the regression analysis revealed significantly larger horns on the LCP relative to the mainland, with the polynomial model explaining a moderate amount of the total variation in horn length (R^2^ = 0.45; Fig. 2). Growth during the first year was higher on the LCP compared to the mainland (18.3 cm ± 1.5 vs 16.4 cm ± 1.6; *t*-statistic 2.68, *p* = 0.05). A similar trend was observed in the second annuli (4.4 cm ± 0.6 vs 3.8 cm ± 3.2; *t*-statistic 1.46, *p* = 0.18).

**Figure 2.**
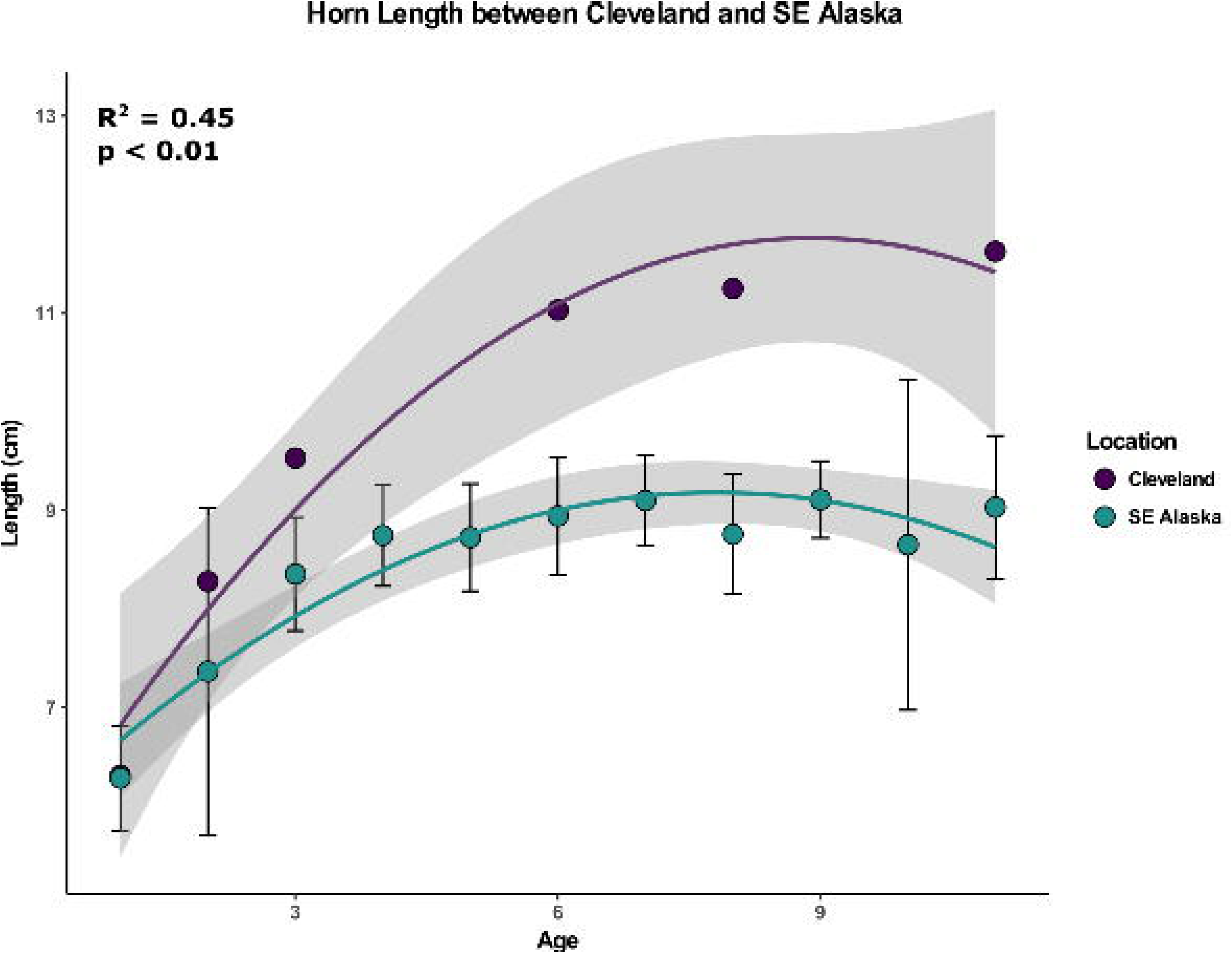
Plot of the polynomial model explaining a total variation in horn length with age between the Cleveland Peninsula and samples from southeast Alaska.

A total of 9 days was spent helicopter sampling pellets, and a large portion of the peninsula was sampled (Fig. S1). We collected a total of 639 pellet groups and extracted 214 tissue samples from our harvested mountain goat tissue repository. We note that preservation in Longmire’s solution resulted in more successful amplification compared to ethanol and dry storage. After data filtering (missing data, duplicates) a total of 468 samples were genotyped; 201 from the Mainland (tissue), 15 from the transition area Bradfield-Stikine (pellet), and 252 from the Cleveland Peninsula (239 pellet, 13 tissue). All 18 loci had less than 50% missing data and were used, with the overall genotyping data set 84% complete (Dryad Accession no. XXXXX). Comparing fecal to the tissue samples on the LCP revealed similar diversity statistics (Table S3).

Estimates of *F*_ST_ between our samples from mainland (GMU 01A, 01B, 01C) and the Cleveland Peninsula, excluding B-S, ranged from 0.02-0.12 with *N*_m_ at approximately 3 migrants per generation (Table S2). Between the four grouped populations on the Peninsula and B-S, pairwise *F*_ST_ ranged from 0.02-0.11 (Table S4), with B-S being the most differentiated. Observed heterozygosity (*H*_O_) for 01A, B, and C ranged from 0.41 to 0.48 (Table 2). Within the groupings on the Cleveland Peninsula, *H*_O_ ranged from 0.23 to 0.34, with B-S at 0.40; thus genetic diversity appears to decrease as mountain goat populations move down the peninsula, with inbreeding coefficients also becoming elevated (Table 2). There was a clear pattern of isolation-by-distance (Mantel *r* = 0.60; p < 0.01; Fig. 3). Our estimates of *N*_e_ ranged from 5-15 individuals (Table 2) for the Lower Cleveland Peninsula using the lowest allele frequency. Estimates of *N*_e_ for the total mainland metapopulation ranged from 36-120 (Table 2). There was no relationship between sample size and estimate of *N*_e_ (*β* = 0.28; *p* = 0.28; Fig. S2).

**Table 1.**
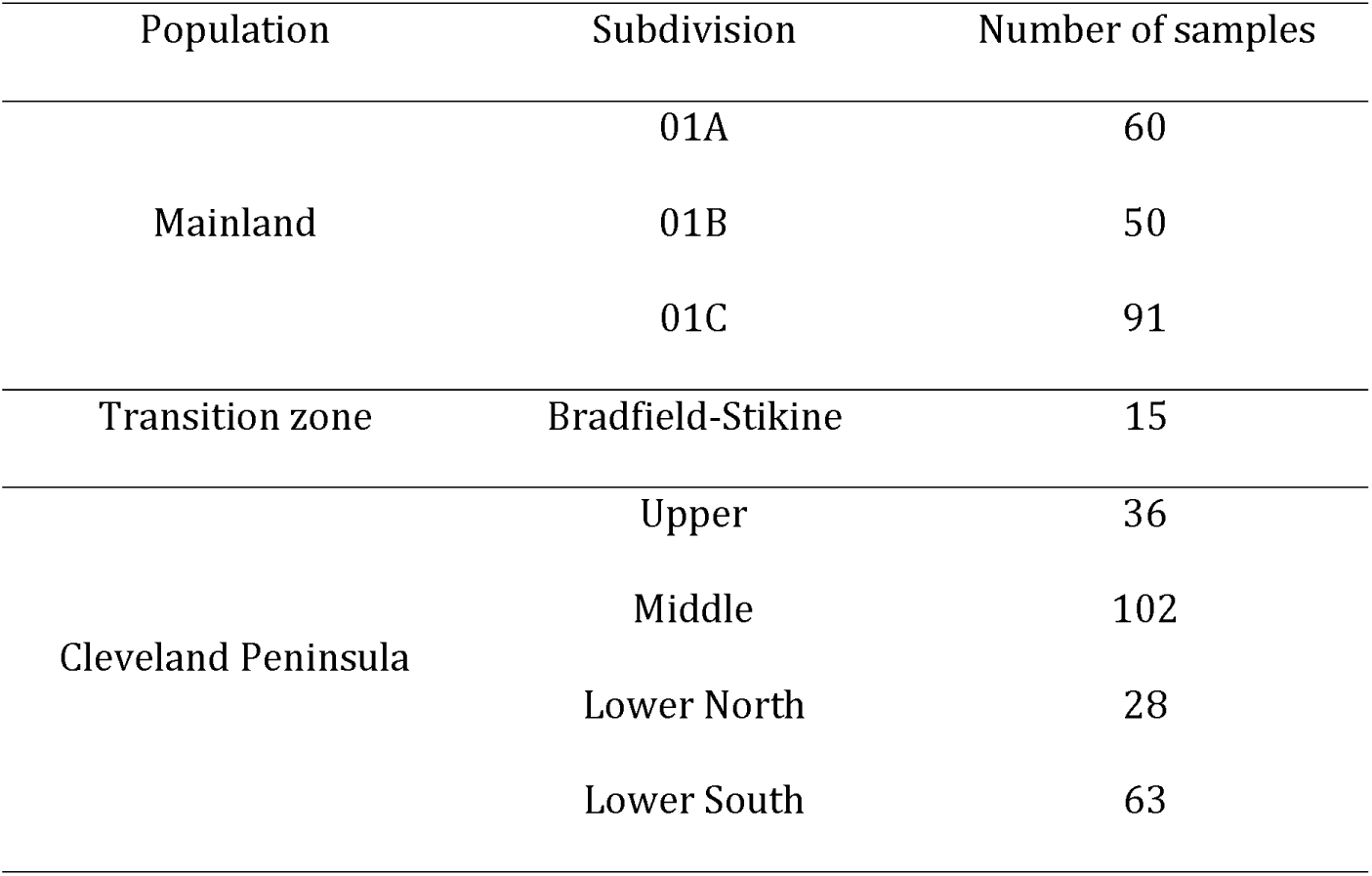
Sample numbers from the Cleveland Peninsula and the mainland used in the genetic analysis. The subdivisions are shown on Fig. 1.

**Table 2.** Diversity statistics for the Mainland the Cleveland Peninsula. The effective population size estimate (*N*_e_) reflects the upper and lower estimates. The subdivisions are shown on Fig. 1.

| Population | Subdivision | No. of Alleles | Observed Heterozygosity | Fixation Index | $N_e$ estimate |
| --- | --- | --- | --- | --- | --- |
| Southeastern Alaska | 01A | $4.8 \pm 0.61$ | $0.41 \pm 0.05$ | $0.10 \pm 0.04$ | 36.0-58.1 |
| | 01B | $5.4 \pm 0.58$ | $0.46 \pm 0.04$ | $0.17 \pm 0.04$ | 81.4-119.4 |
| | 01C | $5.6 \pm 0.54$ | $0.48 \pm 0.04$ | $0.10 \pm 0.03$ | 64.3-95.7 |
| Transition zone | Bradfield-Stikine | $3.1 \pm 0.30$ | $0.40 \pm 0.05$ | $0.08 \pm 0.08$ | 2.0-2.4 |
| Cleveland Peninsula | Upper | $3.9 \pm 0.56$ | $0.34 \pm 0.04$ | $0.30 \pm 0.06$ | 14.7-17.5 |
| | Middle | $4.8 \pm 0.64$ | $0.31 \pm 0.04$ | $0.37 \pm 0.05$ | 27.3-47.7 |
| | Lower North | $4.4 \pm 0.46$ | $0.23 \pm 0.04$ | $0.50 \pm 0.07$ | 7.3-14.9 |
| | Lower South | $4.3 \pm 0.44$ | $0.28 \pm 0.04$ | $0.38 \pm 0.05$ | 4.8-7.9 |

**Figure 3.**
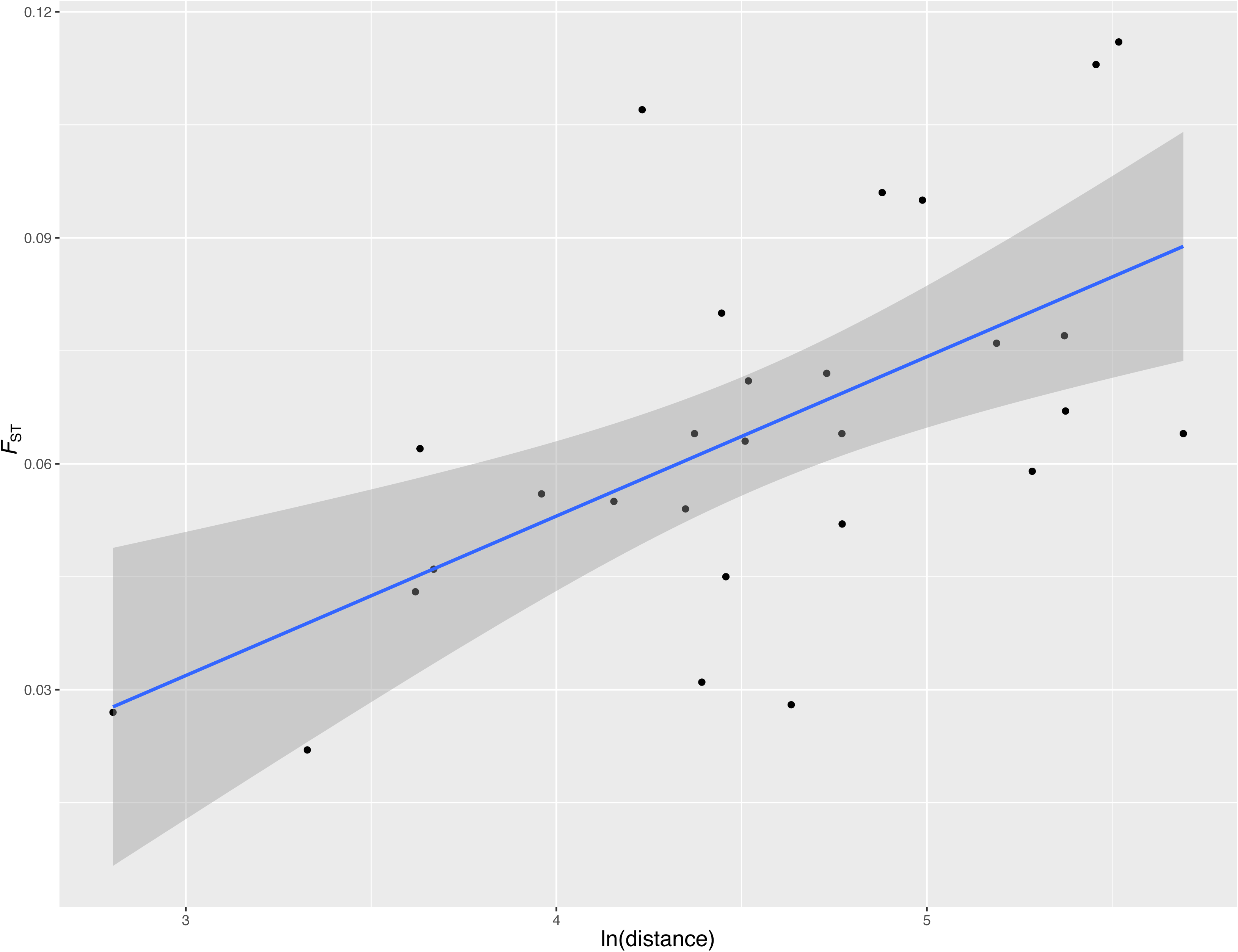
Pattern of isolation-by-distance seen across the study area. Geographic distance is in kilometers and has be log-transformed.

We used the algorithm developed by Evanno et al. (2005) to identify mountain goat group clusters (K). Choice of appropriate number of clusters was informed by Δ(*k*), which identified either *k* = 2 as the optimal number of cluster groups in the genotypic data, which largely divided the Cleveland Peninsula from the mainland (Fig. 4; *k* of 2-5 are also shown). Notably a *k* of 5 is consistent with the isolation-by-distance pattern, which is also seen in the PCA (Fig. S3).

**Figure 4.**
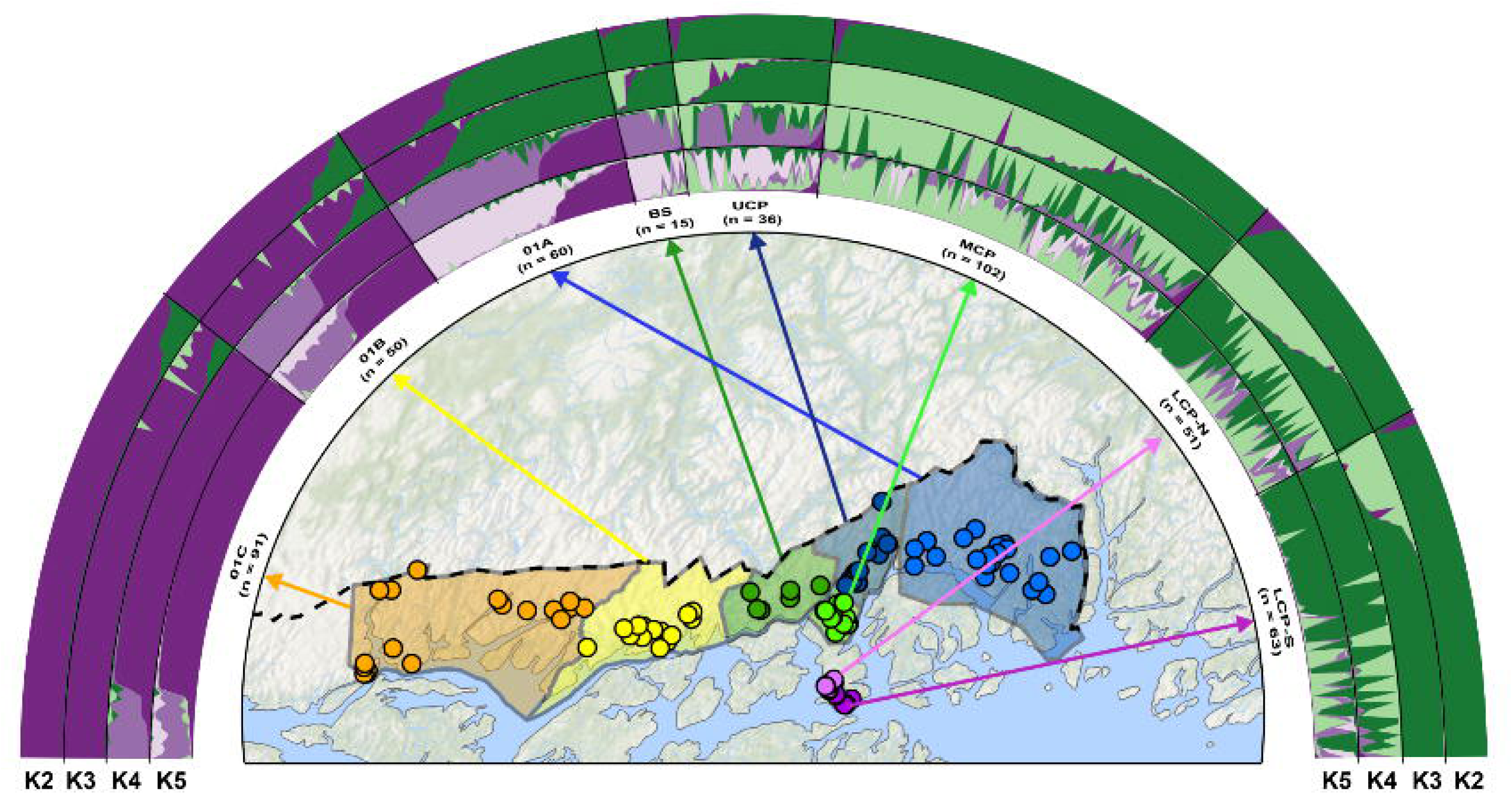
Results of STRUCTURE analysis showing genetic subdivision at *K* 2 through 5. A map showing population assignments is provided; Lower Cleveland Peninsula North (LCP-N), Lower Cleveland Peninsula South (LCP-S), Middle Cleveland Peninsula (MCP), Upper Cleveland Peninsula (UCP) and Bradfield-Stikine (BS) and management units 01A, 01B and 01C.

We sequenced the mitochondrial DNA of 94 samples from the CP (GenBank Accession no. XXXXX). We added 38 samples from southeast Alaska that were available on GenBank. From our 94 mtDNA sequences from the CP we identified 21 unique haplotypes, with the majority found only on the peninsula and not shared with the mainland (Fig. 5)

**Figure 5.**
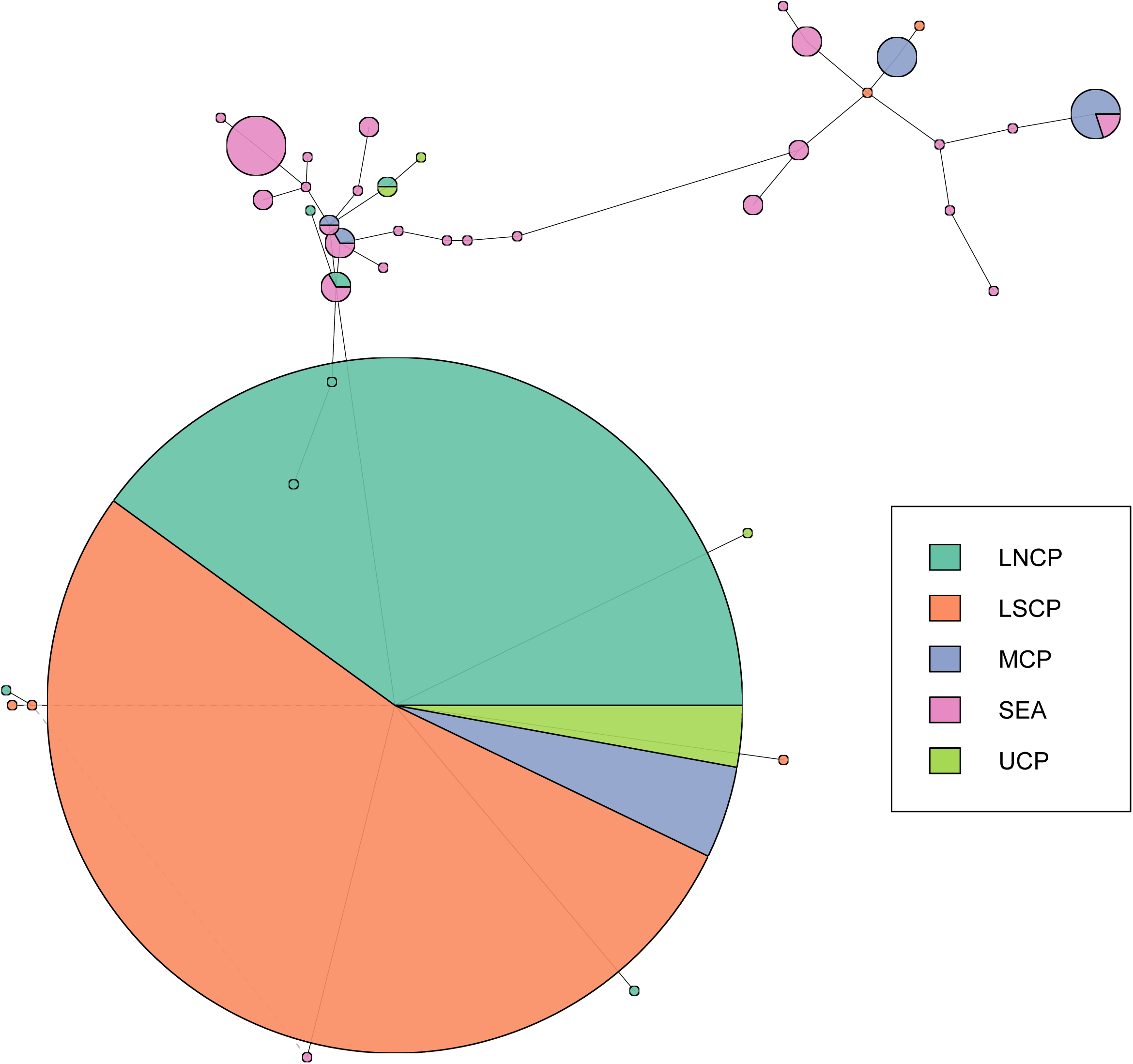
Haplotype network showing difference in mitochondrial sequence between the Cleveland Peninsula (*n* = 95) and samples from the mainland (*n* = 38). The key unique haplotypes unique to the Cleveland Peninsula are highlighted in red.

## Discussion

The results of our analysis suggest that the subpopulation(s) on the Cleveland Peninsula are phenotypically and genetically differentiated from the adjacent GMU situated in southeast Alaska. Metrics of diversity suggest the Cleveland Peninsula is isolated with limited gene flow and is thus at risk for inbreeding depression and increased extinction risk (Caballero et al. 2017). Populations on the peninsula do appear interconnected, constituting a meta-population, with a pattern consistent with stepping-stone model, meaning decreasing diversity as you move down the peninsula due to decreased migration. Collectively, this suggests the Cleveland Peninsula is largely isolated from the mainland and demographically distinct. How these data and patterns can inform actual management, however, is more nuanced.

### Interpreting genetic parameters for management

A gap has emerged between genetic studies and on-the-ground conservation and management needs (Shafer et al. 2015, Taylor et al. 2017). While there are a plethora of reasons for this gap (see Taylor et al. 2017, Britt et al. 2018, Mair et al. 2018), part of the problem stems from the abstract and relative nature of many genetic parameters making them difficult to interpret and implement. General convention describes that *F*_ST_ < 0.05 constitutes little genetic difference, *F*_ST_ = 0.05-0.15 as moderate genetic difference, *F*_ST_ = 0.15-0.25 as large genetic differentiation, and *F*_ST_ > 0.25 as significant genetic difference (Hartl and Clark 1997). Thus, the Cleveland Peninsula population can be considered moderately differentiated from the mainland. We cannot, however, distinguish whether the population is recently isolated and there is currently no gene flow, meaning signatures of *N*_m_ simply stem from recent common ancestry, or the CP mountain goats have been isolated for a long-time with periodic migration. The large number of individuals containing the CP unique mtDNA haplotypes would support the latter, further supporting the demographic uniqueness of this population.

Not only do the data support a demographically distinct population on the Cleveland Peninsula, but one that is vulnerable to environmental and demographic perturbations. The estimate *N*_e_ is a key population genetic parameter described by Wright (1931) as the theoretical number of individuals in an ideal population that would lose genetic variation at the same rate as the observed population (i.e. the census size or *N*_C_). Interpreting this parameter is difficult, as the approach reflects the parental generation’s *N*_e_, but can be impacted by changes further in the past (Waples 2005). Efforts to link *N*_e_ to *N*_C_ are desirable as they allow for going from a genetic parameter to the number of individuals on the landscape; however, *N*_e_/*N*_C_ ratios often span multiple orders of magnitude leading to high uncertainty (Palstra and Fraser 2012). Ortego et al. (2011) found the ratio of *N*_e_/*N*_C_ between 0.4 and 0.6 in mountain goat population found on Caw Ridge, Alberta. This seems reasonable for the lower Cleveland Peninsula given the *N*_e_ of less than 15 and aerial survey estimates from the 1980’s of 50-70 individuals, but the key take-home is reduced diversity on the LCP and UCP relative to the mainland.

The small effective population size and high inbreeding coefficient on the lower Cleveland Peninsula is an indication of a lack of genetic variability and may lead to reduced fitness of individuals, an increase of inbreeding, and further reduction in population size if not managed accordingly (Armbruster and Reed 2005). Habitat on the lower Cleveland Peninsula is atypical for mountain goats (Smith and Raedeke 1982), and it is unlikely sufficient suitable habitat and resources exist to accommodate more individuals. Without gene flow from the mainland it is reasonable to predict genetic diversity will continue to decrease if population numbers do not change. Low *N*_e_ combined with minimal migration to the mainland equates to reduced adaptive potential, making the population more susceptible to environmental and demographic perturbations than their mainland counterparts.

## MANAGEMENT IMPLICATIONS

Given the geographic isolation, phenotypic differences, and the genetic distinctiveness reported here, the current harvest moratoria and separate hunting is warranted on the LCP (e.g. Moritz 1994, Fraser and Bernatchez 2001). These data suggest a sustainable harvest is unlikely on the LCP given the genetic parameters and low population numbers; if the population demographically recovers, any future harvest should be limited to adult males (Gonzalez-Voyer et al. 2003) and consideration should be given to alternating harvest years given the high extirpation risk (Hamel et al. 2006). The UCP and MCP are clearly more connected to the mainland, but it too is demographically distinct, and likely warrants a separated hunting unit status from the LCP. Given that the MCP and UCP are hard-to-access areas, a high impact of harvest seems unlikely To prevent further loss of genetic variability in the region, translocating individuals from southeast Alaska to the Cleveland Peninsula is a management option. While this could impact the unique and desirable phenotype on the peninsula (Fig. 3), the translocation of even a few individuals carrying novel alleles could be enough to genetically augment this population if needed (Hufbauer et al. 2015). That said, given the atypical habitat on the lower Cleveland Peninsula it is not clear that the region could even support the number of individuals that are desirable for a sustained harvest (see Hamel et al. 2006), and thus translocations should be considered a last resort.

## Supporting information

## ACKNOWLEDGEMENTS

We thank Sarah Haworth and Jesse Wolf help with sequencing. Susannah Woodruff provided valuable feedback on this manuscript. JB was supported by an EcoCanada Internship; MB by a NSERC undergraduate student research award and DM by an Ontario Graduate Scholarship. The project was supported by NSERC-DG, CFI-JELF and Compute Canada grants to ABAS; US Federal Aid grants to KSW and ABAS.

