## Supplementary material for "Extensive field-sampling reveals the uniqueness of a trophy mountain goat population"

**SUPPLEMENTAL FIGURES AND TABLES**

Figure S1. Map of all pellet samples collected in this study. Study site is in southeast Alaska.

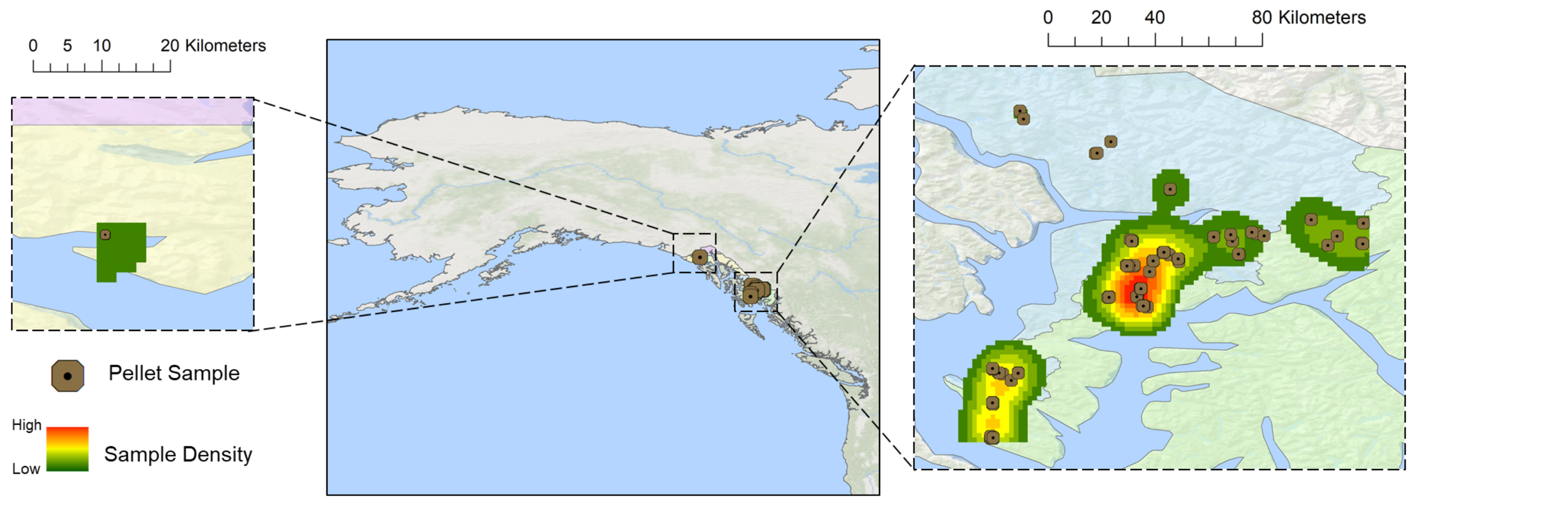

Figure S2. Relationship between sample size and lowest *N*_E_ estimate (see Table 2).

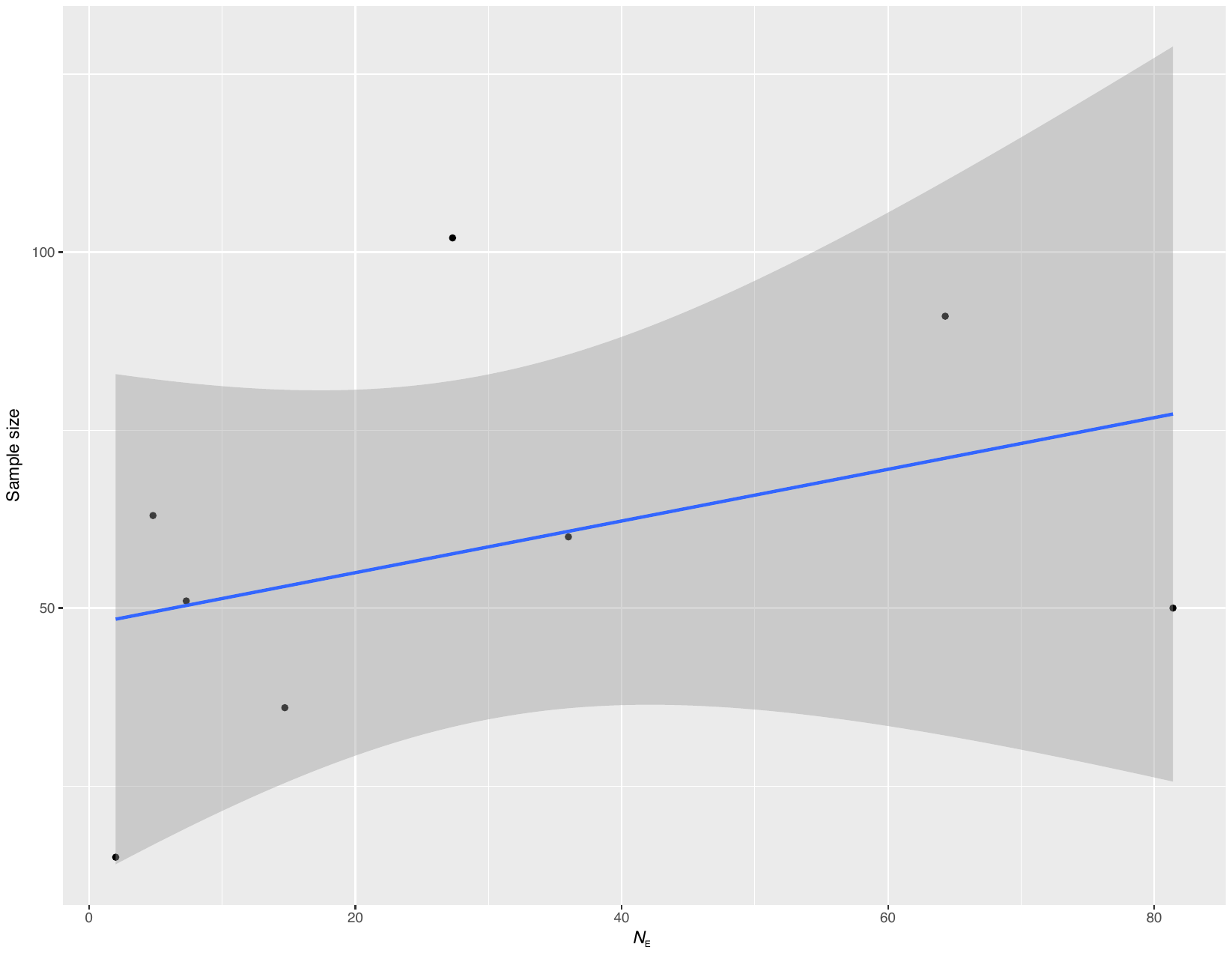

Figure S3. PCA of final genetic data set.

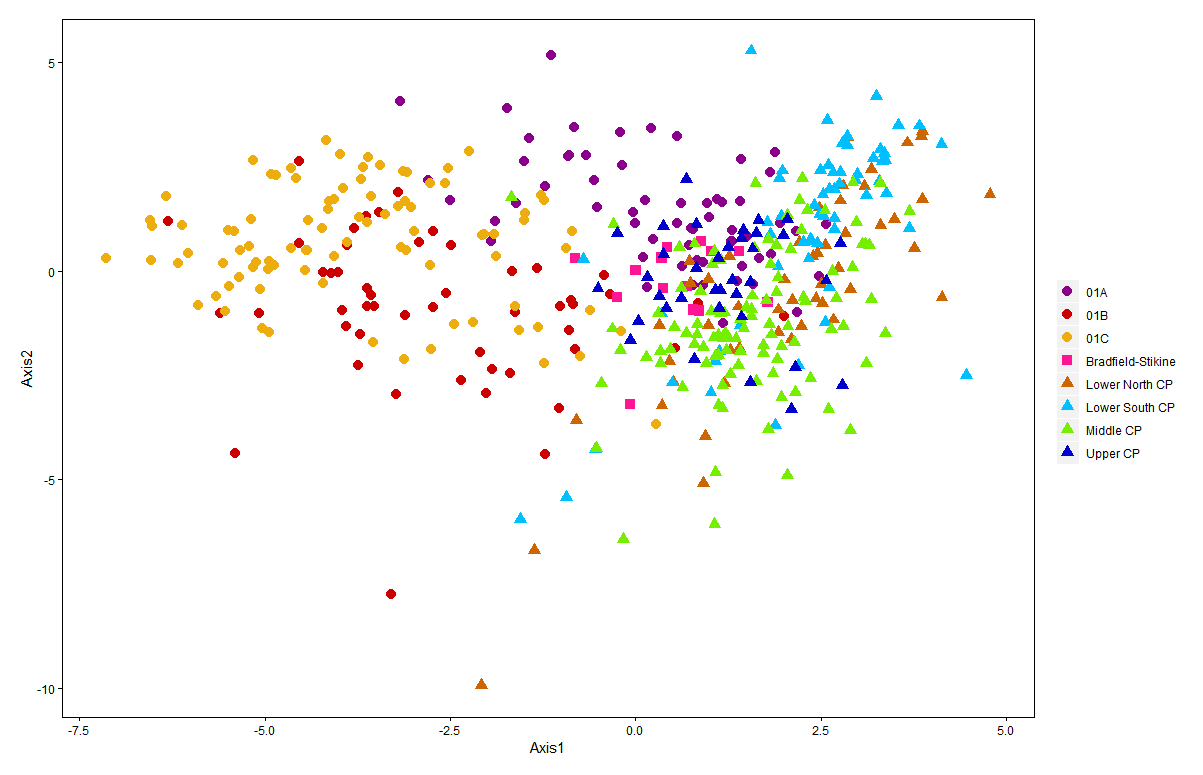

Table S1. Microsatellite loci and pools used to genotype mountain goats in this study.

|  | Locus |
| --- | --- |
| Pool 1 | MAF36 |
|  | TGLA122 |
|  | OARJMP58 |
|  | TGLA10 |
|  | ILSTS058 |
|  | RT9 |
| Pool 2 | BR3510 |
|  | MCM152 |
|  | ARO28 |
|  | OARCP26 |
|  | BM1225 |
|  | OARJMP29 |
| Pool 3 | OARHH62 |
|  | MAF64 |
|  | HUJ1177 |
|  | OARHH35 |
|  | RT27 |
|  | HUH616 |

Table S2. Pairwise F_ST_ (lower matrix) and migration rate per generation (upper matrix). Populations are: Lower Cleveland Peninsula North (LCP-N), Lower Cleveland Peninsula South (LCP-S), Middle Cleveland Peninsula (MCP), Upper Cleveland Peninsula (UCP); Bradfield-Stikine (BS) and GMUs 01A, 01B, and 01C – see Fig. 1.

|  | 01A | 01B | 01C | B-S | LCP-N | LCP-S | MCP | UCP |
| --- | --- | --- | --- | --- | --- | --- | --- | --- |
| 01A | - | 4.0 | 3.7 | 4.5 | 3.3 | 3.7 | 5.3 | 7.7 |
| 01B | 0.059 | - | 8.5 | 3.6 | 2.3 | 2.4 | 3.2 | 3.6 |
| 01C | 0.064 | 0.028 | - | 3.1 | 2.0 | 1.9 | 3.0 | 3.5 |
| B-S | 0.052 | 0.064 | 0.076 | - | 2.1 | 2.9 | 3.8 | 5.2 |
| LCP-N | 0.071 | 0.096 | 0.113 | 0.107 | - | 9.0 | 5.5 | 4.3 |
| LCP-S | 0.063 | 0.095 | 0.116 | 0.080 | 0.027 | - | 4.2 | 4.4 |
| MCP | 0.045 | 0.072 | 0.077 | 0.062 | 0.043 | 0.056 | - | 11.2 |
| UCP | 0.031 | 0.064 | 0.067 | 0.046 | 0.055 | 0.054 | 0.022 | - |

Table S3. Diversity statistics between tissue and non-invasive samples on from the Lower Cleveland Peninsula.

| Population | No. of samples | No. of alleles | Observed heterozygosity |
| --- | --- | --- | --- |
| Tissue | 13 | 2.444 ± 0.271 | 0.286 ± 0.065 |
| Non-invasive | 101 | 4.778 ± 0.461 | 0.245 ± 0.034 |

Table S4. Pairwise F_ST_ (lower matrix) and migration rate per generation (upper matrix) values between the subpopulations on the Cleveland Peninsula - Lower Cleveland Peninsula North (LCP-N), Lower Cleveland Peninsula South (LCP-S), Middle Cleveland Peninsula (MCP), Upper Cleveland Peninsula (UCP), and Bradfield-Stikine (B-S) – see Fig. 1.

|  | B-S | UCP | MCP | LCP-N | LCP-S |
| --- | --- | --- | --- | --- | --- |
| B-S | - | 5.2 | 3.8 | 2.1 | 2.9 |
| UCP | 0.046 | - | 11.2 | 4.3 | 4.4 |
| MCP | 0.062 | 0.022 | - | 5.5 | 4.2 |
| LCP-N | 0.107 | 0.055 | 0.043 | - | 9.0 |
| LCP-S | 0.080 | 0.054 | 0.056 | 0.027 | - |
